# Predicting nucleation sites in chemotaxing *Dictyostelium discoideum*

**DOI:** 10.1101/564963

**Authors:** E.O. Asante-Asamani, Devarshi Rawal, Zully Santiago, Derrick Brazill, John Loustau

**Affiliations:** Department of Mathematics and Statistics, Hunter College, New York, NY USA; Department of Biological Sciences, Hunter College and the Graduate Center, CUNY, New York, NY USA; Department of Biological Sciences, Hunter College, CUNY, New York, NY USA

**Author notes:** These authors also contributed equally to this work (E.O. Asante-Asamani, John Loustau).

## Abstract

Blebs, pressure driven protrusions of the plasma membrane, facilitate the movement of cell such as the soil amoeba *Dictyostelium discoideum* in a three dimensional environment. The goal of the article is to develop a means to predict nucleation sites. We accomplish this through an energy functional that includes the influence of cell membrane geometry (membrane curvature and tension), membrane-cortex linking protein lengths as well as local pressure differentials. We apply the resulting functional to the parameterized microscopy images of chemotaxing *Dictyostelium* cells. By restricting the functional to the cell boundary influenced by the cyclic AMP (cAMP) chemo-attractant (the cell anterior), we find that the next nucleation site ranks high in the top 10 energy values. More specifically, if we look only at the boundary segment defined by the extent of the expected bleb, then 96.8% of the highest energy sites identify the nucleation.

**Author summary:** This work concerns the prediction of nucleation sites in the soil amoeba-like *Dictyostelium discoideum*. We define a real valued functional combining input from cortex and membrane geometry such as membrane curvature and tension, cortex to membrane separation and local pressure differences. We show that the functional may be used to predict the location of bleb nucleation. In the region influenced by the cAMP gradient (the cell anterior), the next blebbing site lies in the ten highest energy functional values 70% of the time. The correctness increases to 96.8% provided we restrict attention to the segment in the general location of the next bleb. We verify these claims through the observation of microscopy images. The images are sequential at 1.66 and 0.8 seconds per image. We first identify the earliest sign of the bleb. We then use several observational factors to identify the nucleation site and estimate the corresponding location in the prior image.

## Introduction

For *Dictyostelium discoideum* (*D. discoideum*) as well as various mammalian cell types, the means of motility changes depending on varying conditions. At least since Yoshida and Soldati [37], we have known that *D. discoideum* cells in three dimensional environments often use blebs as a means of cell extension. These are pressure driven blister-like protrusions of the cell membrane. Further, the greater the pressure from the environment, the greater the reliance on blebs during migration. In humans, well-known cases of cells using blebbing to migrate include embryonic organ development and cancer cell migration ([6], [33], [15], [26], [2], [13]).

*D. discoideum* is a species of soil-living amoeba-like organism. It is a eukaryotic organism whose motility and shape is controlled predominantly by intricate actin based structures in the cytoplasm. In turn, the cell cytoplasm is encased by a plasma membrane, a 4-5nm thick semi-permeable lipid bilayer [38]. Beneath this membrane is the cell cortex, an assembly of thin cross-linked actin filaments held together firmly by cross linking proteins such as Filamin, *α*-Actinin, Fimbrin and Fascin. The cortex thickness is several hundred nanometers [1]. The cortex plays a major role in maintaining cell shape and the formation of motility structures. Its contractile capabilities are due to the presence of Myosin II in the network. The membrane is attached to the cortex by trans membrane proteins such as Talin [9]. In addition to the cortex, the cytoplasm includes distinct actin-based structures referred to as the cytoskeleton. The fluid part of the cytoplasm (cytosol) contains water, ions and dissolved molecules.

Blebs and blebbing have been approached from several viewpoints. These studies may describe blebs as geometric objects (Euclidean) [36], the result of physical force (pressure) [7], [5], [17], the result of cortical tension (related to pressure) [32], the result of fluid dynamics [30], [29] and forces on the boundary of a smooth manifold (differential geometry) [34], [39], [21] and [9].

By and large, the question of the cause of bleb nucleation has not been resolved. Most researchers agree that blebs are formed by the action of pressurized cytosol on the plasma membrane [11], [37], [6]. This pressure is the result of local contraction of the actin cortex by Myosin II motor proteins at the posterior end. This causes a flow of cytosol towards the anterior. What differs among the theories of nucleation is the mechanism by which the plasma membrane detaches. According to Charras *et al*. [5], the membrane-cortex detachment is the result of the pressure differential between the posterior and anterior. The pressure differential dislodges the membrane from the cortex. This point of view is supported by Pullarkat [24] and Young and Mitran [38]. Also supporting this point of view, Collier *et al* [9] show that Talin is concentrated at the posterior where there is little blebbing while at the anterior the Talin is sparsely distributed in *D*. *discoideum*. Alternatively, Paluch *et al* [22] and Keller and Eggli [19] suggest that membrane-cortex detachment is enhanced by local degradation of the actin cortex. At these locations, the membrane no longer has the support of the cortex. The force of the flow is then capable of breaking the adhesive bonds and detaches the membrane. Interestingly, Charras *et al* [7] also provide evidence for the local cortex degradation through the action of Myosin II. They are seemingly on both sides of the debate.

In this study, we approach this issue from a different point of view. We begin with the solid foundation of our geometric platform [27]. When under environmental pressure, *D. discoideum* cells are largely two dimensional. Hence, like our predecessors we use two dimensional geometric constructs. With this foundation, we have developed an energy functional and used it to predict bleb nucleation sites. This functional is a modified version of the one presented in [9] and [34] or alternatively the one in [21]. Both are derived from an interactive graphics application developed in [18] from work originated by Helfrich to model the shape of red blood cells [16], [14].

According to [18], any shape has an energy cost. This idea goes back to Euler-Lagrange that provides the solution to the related calculus of variation problem [12]. Any unconstrained shape will seek to transform into a shape of lower energy cost. We expand this idea and propose that blebs occur at locally high energy locations. Subsequently, the convex, circular bleb has nearly constant energy. Hence, there are no longer high energy spikes and the cost of maintaining the shape is lowered.

Secondly, we focus our attention to the area of the cell affected by the gradient of cAMP, the area of blebbing activity. In our setting, this is roughly half the cell boundary. Then we ask whether the energy functional can identify the specific nucleation site.

For chemotaxing and confined *D. discoideum* cells under 0.7% agarose, we identify newly formed blebs and use various observational methods to identify the nucleation site. These cells generally exhibit high energy profiles in the blebbing region. For cases that we can infer the nucleation site in the image prior to the bleb, this location coincides with the local high energy location identified by the functional in over 96% of the observed cases. Furthermore, when applying the functional to the general region affected by cAMP, our energy functional located the next nucleation site 40% of the time as the first or second top energy value.This finding is comparable to results reported in [9].

This project has used differential geometric and local pressure differential to predict bleb initiation site. In addition, in the process of locating nucleation sites, we have noted frequent evidence of cortex degradation just prior to blebbing. This validates the point of view that cortex ruptures initiates belb nucleation. We have tabulated these events below. Subsequent biological research may be able to clarify the interplay between the differential geometry and the biology in bleb site selection and explain the observed early cortex disassembly.

This article is organized as follows. In the Methods and Materials section, we review the cell preparation and microscopy technology. We state the geometric procedures underlying this work and the basic boundary energy functional to be used. At that time we list the separate energy functional components. In Results and Discussion we derive the discrete form of the energy functional with all parameters resolved. Then we carry out a study to test the functional against ROI observation in predicting bleb nucleation sites. Finally, we compare the individual components of the functional against the full functional. In the Conclusion we summarize what we have done and consider possible future biological work and more sophisticated numerical processing.

## Materials and Methods

### Strain and culture conditions

All *D. discoideum* cells were grown axenically in shaking culture in HL5 nutrient medium with glucose (ForMedium) supplemented with 100 *µ*g/mL penicillin and 100 *µ*g/mL streptomycin (Amresco) at 150 rpm at 22°C with 4-20ug/mL G418 (Geneticin) for LifeAct-GFP expressing Ax2 wild-type cells [27]. Cells were starved for cAMP competency on filter pads using the method described in [27].

### cAMP under agarose assays

cAMP under agarose assays were used as described in [26]. Cells crawled under a 0.7% Omnipur@ agarose (EMD Millipore) gel that was laced or not laced with 1mg/mL of 70,000 MW Rhodamine B isothiocyanate-Dextran (Sigma-Aldrich).

### Image Acquisition

All imaging data were collected using a Leica DMI-4000B inverted microscope (Leica Microsystems Inc.) mounted on a TMC isolation platform (Technical Manufacturing Corporation) with a Yokogawa CSU 10 spinning disc head and Hamamatsu C9100-13 ImagEM EMCCD camera (Perkin Elmer) with diode lasers of 491 nm, 561nm, and 638nm (Spectra Services Inc.) [24] using the same Volocity software and parameters as described in [26]. To summarize, LifeAct-GFP and RITC-Dextran were excited using the 491nm and 561nm lasers, respectively. Cells were imaged for 30 seconds using either 80x magnification (40x/1.25-0.75 oil objective with a 2x C-mount) or 100x (100x/1.44 oil immersion objective). Data collected using both GFP and RITC channels resulted in one frame per 1.66 seconds where data collected using only GFP resulted in one frame per 0.800 seconds. ImageJ was used to adjust the brightness and contrast of the images, which were then imported into our *Mathematica* based geometric system.

### Digitizing Microscopy Images

We use our own system to render photographic images as objects in Cartesian space. In [27], we detail the system components and provide reason that this is appropriate for this setting.

in particular, we apply our procedures to the microscopy output to produce a cubic B-spline representation of the cell. We use this view of the cell or alternatively the view produced by a sequence of equally spaced (arc length) points on the B-spline. The point spacing is approximately 100nm or 0.5 pixel units. (Note: 5 pixels is approximately 1 *µ*m.) We refer to these points collectively as the *EquList*. This is the foundation for the discrete form energy functional.

### An Energy Functional

We use a membrane energy functional related to the one introduced by T. Bretschneider in [34] and [9]. This functional is a modified version of one used in a computer graphics application [18], which in turn is modified from one that arose in modeling red blood cells [14]. Our energy functional (1) is identical to the one defined in [9] save for the last term in the integrand.

In another setting, something similar was used to derive a model for blebbing [21]. In this paper and [9], the energy functional is used to identify locations likely to bleb. In this section, we use a third version of the functional to compute energy values. Our changes are discussed below. Our purpose is the same.

Note that the fourth term of the integrand in (1) is not energy. Hence, the integrand is not energy and the integral is not total energy. We denote it as *pE*_*total*_ for pseudo-energy, and refer to it as energy as the alternative is cumbersome. Moreover, others have called a similar expression energy [9].

At this time we lay out the functional and present the separate elements. We begin with the notation. Points on the membrane are denoted 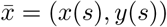, points on the cortex by 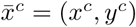, while link lengths are inferred by the distance between the curves. In the discrete model this reduces to the distance between corresponding points in the respective point lists.

The total energy of the complex is expressed as the point-wise energy integrated (summed) along the cell boundary.

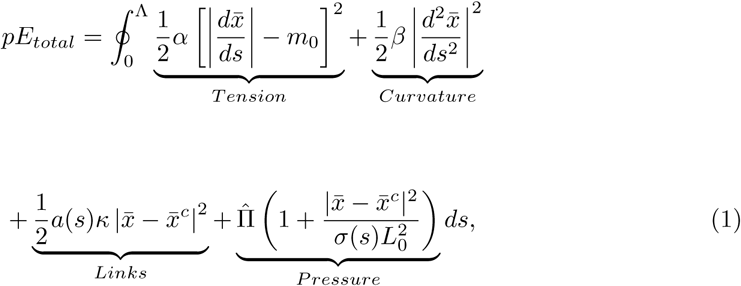

where Λ denotes the total boundary length. The individual terms are described as follows.

- The first two terms refer to the membrane mechanics with membrane stiffness parameter, *α*, and bending rigidity parameter, *β*. The term *m*_0_ is the resting membrane length. The first term is the energy derived from the *Tension*, the second term is the energy required to bend the membrane [16]. Parameter estimates *α, β* and *m*_0_ are provided in *Results and Discussion*.
- The third term is energy associated to link tension. It is determined by the linking proteins modeled as a linear spring between corresponding cortex/membrane points with spring constants as parameters. Setting, 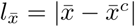, important values of 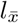 are 0.03*µ*m, the minimal link length, *l*_0_ = 0.056*µ*m, the breaking length. The breaking length is in effect the maximal length of a link. Finally,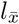 at rest will be about 0.04µm. The spring constant *κ* is resolved below.
- The last item is the hydrostatic pressure. It is not formatted as energy. The parameters are *l*_0_, the breaking length of the links, *σ*(*s*) = *σ* the local multiplier required to scale the data, and 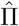 the ambient pressure differential. Values of *σ*(*s*) are computed so that the ratio 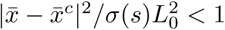 The value of this term reflects the activity of Myosin II. Local pressure raises as a result of cortex contraction by Myosin II. For the most part, these small increases are canceled by other small changes in the cortex/membrane. But this process is not perfectly efficient. Hence, we accommodate for small local variations in pressure. In [4], this effect is identified in cancer cells. In [2], there is a similar report for zebra fish cells. Since we cannot measure pressure directly, we settle for an alternative means. If the actin cortex is contracted, we expect increased separation from the membrane. As a result, the linking protein is stretched. Hence, we use changes in the linking protein lengths as a replacement for pressure changes caused by Myosin II contractions.

We caution the reader that the first term of the integrand of (1) is not a constant as it might appear. Rather, the statement in (1) represents the energy at the initial values. Upon initialization, the membrane seeks a position that minimizes the functional. As relaxation occurs, the membrane parameterization is no longer arc length. See also [28].

We may use the integrand in (1) to define pointwise or local energy.

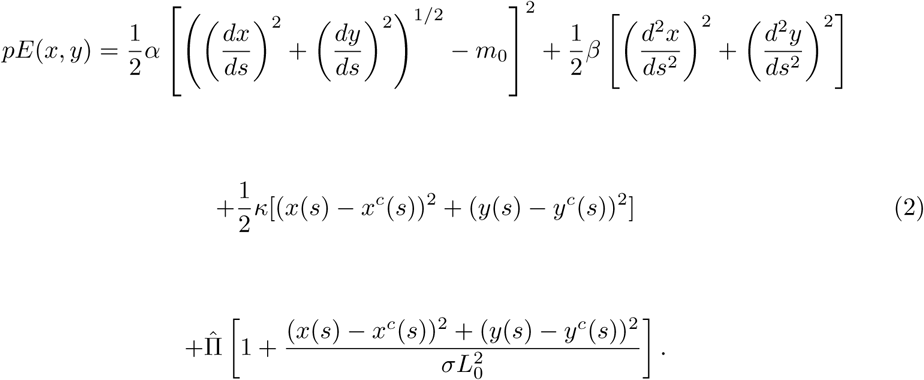

## Results and Discussion

### The Discrete Energy Functional

In order to use the local functional (2) we needed to know the location of the membrane. We knew the cortex as a cubic B-spline or via the EquList 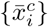 of equally spaced cortex points. Furthermore, we knew that the membrane is so close to the cortex that it is nearly indistinguishable in a microscopy image.

We began by defining the membrane via a list of points 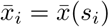 where each point has distance *l*_*i*_ from 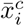 along a ray initiating at an interior point of the cell (usually the cell centroid). We identified this distance with the length of the linking protein joining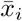 and 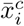 The issue now was to determine the *l*_*i*_.

We proceeded with the assumption that the cortex and membrane are nearly identical, or equivalently, the *l*_*i*_ are very small. Consequently, we supposed that the list 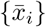 is nearly equally spaced. Furthermore, as we did not know the curve representing the membrane, we used finite differences, specifically central differences, for the derivatives in (2). Next, we restated the local energy functional in discrete form.

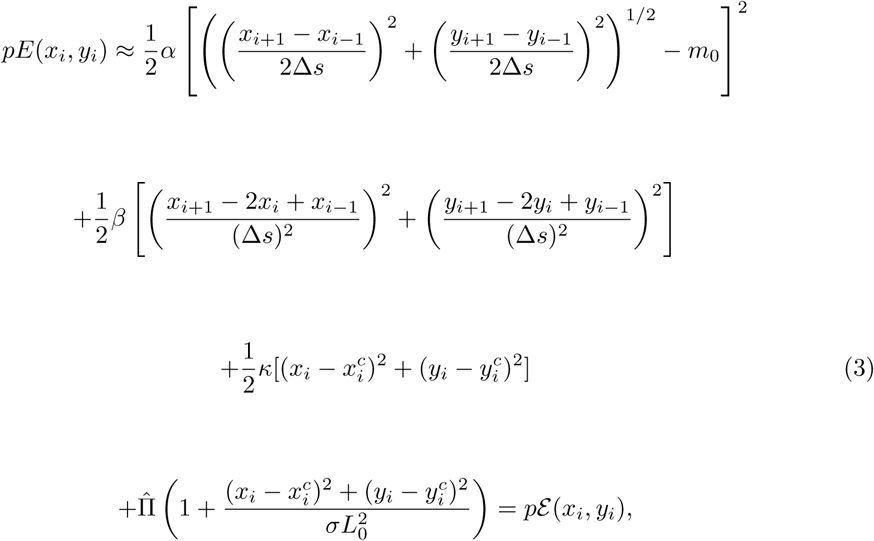

where Δ*s* is the designated fixed distance between points on the membrane list. In turn, we defined the discrete total energy. This is in effect the distance between points on the cortex list.

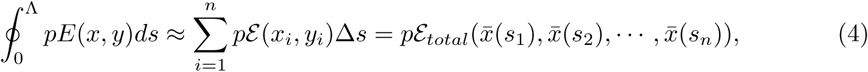

a function of *n* variables, where *n* is the length of the EquList.

### **Resolving the** *l*_*i*_

It remained to determine the *l*_*i*_, equivalently the 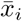 When that was done, we knew *pε*_*total*_ and each 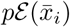.

First we set the parameters for (3) as follows:

- *α* = 17.0 pN/*µ*M,
- *β* = 0.14 pN/*µ*M,
- *κ* = 10.0 pN/*µ*M,
- II = 81 pa.

These values were found in [34].

We resolved the *n* unknowns by arguing that the complex relaxes from an initial state and seeks a stable state. At this state, the energy attains a local minimum, so that any small variation in any *l*_*i*_ increases the energy.

We took an initial value *l*_*i*_ = 0.03*µ*M, the minimal distance between the membrane and cortex. We proceeded with minimization via a gradient directed search [20] and arrived at a local minimum for *pε*_*total*_. We accepted the *l*_*i*_ and 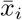 for this state. This done, we could evaluate *pε* (*x*_*i*_, *y*_*i*_) for each 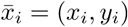.

### Energy and Bleb Nucleation

We began by reviewing how to locate the bleb nucleation site. For instance, consider Fig 1A. Here there are two microscopy images of a cell taken at successive times, just before and just after a bleb occurs. The images, Fig 1A1 and Fig 1A3 show prior and after views of the membrane and the images Fig 1A2 and Fig 1A4 show the prior and after images of the cortex. While there is clear evidence of membrane/cortex separation in the right hand images, A3 and A4, there is no sign of this separation in A1 and A2, the area of the expected bleb.

**Fig 1.**
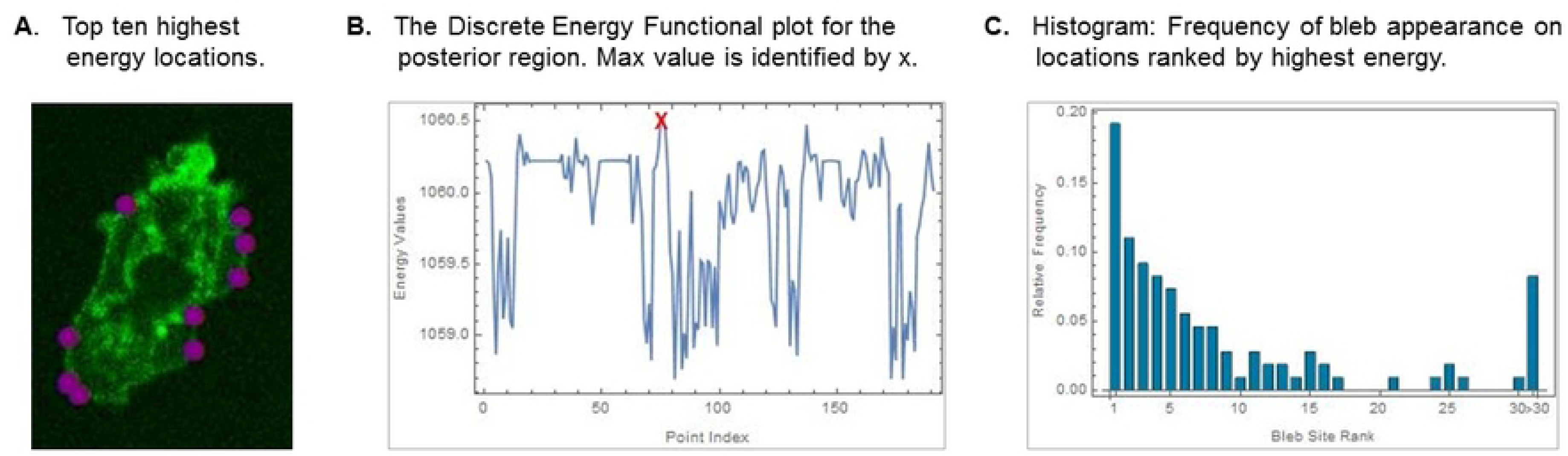
Identifying The Nucleation Site as ROI. (A) Bleb: Pre and Post Images. Pre Bleb images of the cell are shown in A1 and A2. A1 is the membrane image and A2 is the corresponding cortex image. The post bleb images are A3, membrane and A4, cortex. The arrows on A3 and A4 identify the membrane-separation associated to the bleb. Note there is no separation in A1 and A2. (B) Finding bleb features in order to infer nucleation in the pre bleb image. Images B1 and B2 show the cortex just after formation of the bleb. In B1 we identify the bleb shoulder points *a* and *b*, the furthest extent of the bleb is denoted by *c*, the cortex gap is noted *d*. In B2 we have identified GFP bundles on either side of the gap, These features are denoted *α* and *β*. (C) Identifying the post bleb features in the pre bleb image. The shoulder points corresponding to *a* and *b* are identified as *a′* and *b′*. *α′* and *β′* identify the GFP bundles in the prior state. The point *d′* is the predecessor to *d* in A. (D) Plot of discrete energy functional between ′ and *b′*. The maximum value is identified by *x*. This point and its near neighbors form an isolated cluster of high energy values. (E) The maximal energy value is identified on the cell boundary by a purple arrow. It corresponds to *d′* in C.

**Fig 2.**
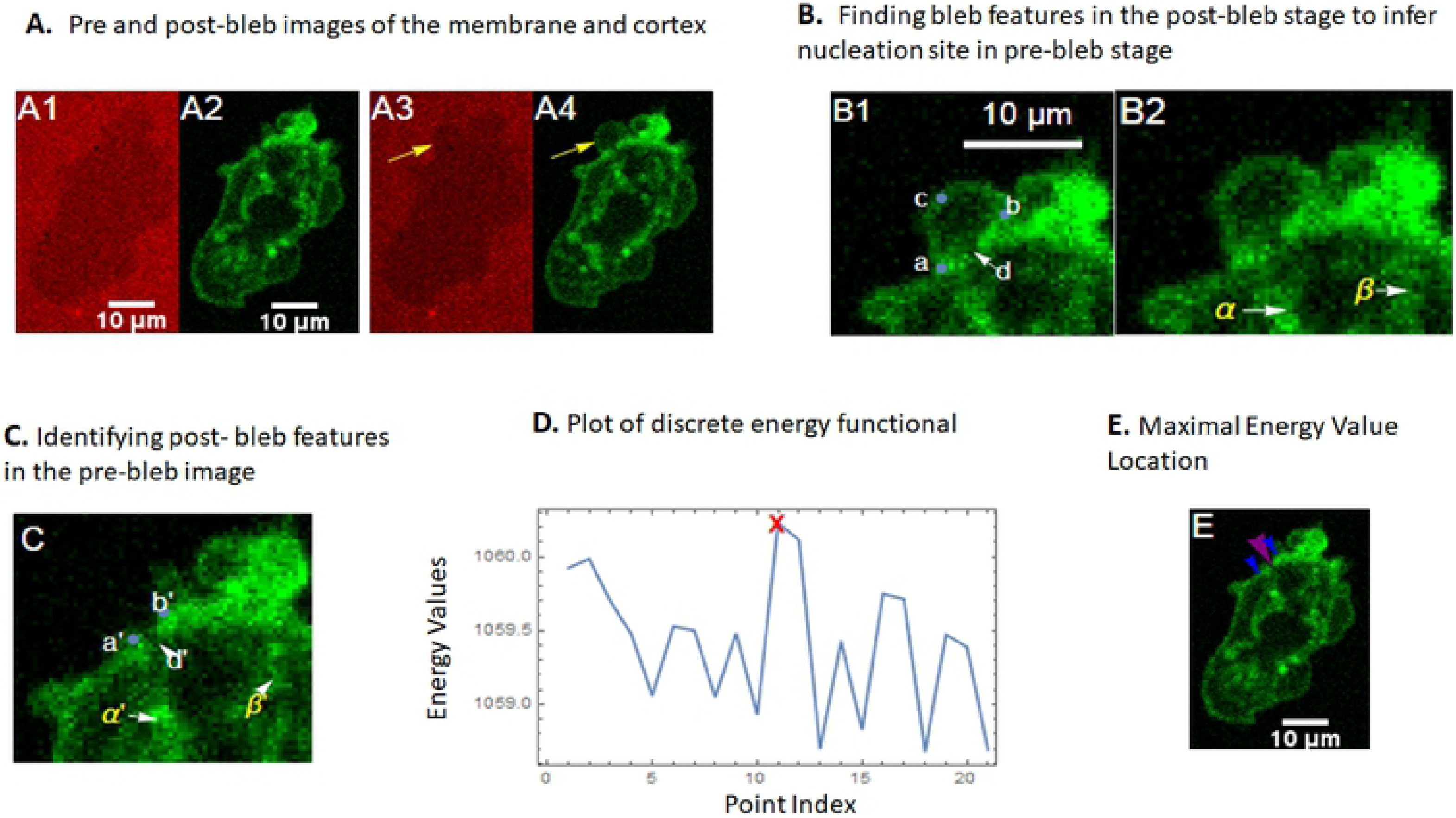
High Energy Locations. (A) The ten highest energy locations for the cell that is about to bleb. Locations are identified with a light blue dot. Note the five high energy locations on the right opposite the pending bleb, shadowed from the cAMP affected area. The region is the posterior where Myosin II activity is known to be clustered. The scale is the same as in Fig 1A. (B) The energy plot for the posterior region showing regional high energy. (C) Histogram showing how the bleb site ranks among the 30 high energy locations on the cAMP affected face. This data is based on 20 cells and 109 blebs. It is statistically significant with 95% confidence.

Clues to the location of the nucleation point can be found in the bleb. There are four features to note. First, we looked for the furthest or maximal extent of the bleb. This is the point furthest from the actin scar or degrading cortex. It is also the oldest part of the bleb with the most developed cortex. We expect the nucleation point to be opposite the maximal extent. This indicator is present even when there are multiple nucleations or modifications of the bleb caused by cortex geometry or neighboring blebs. Hence, it was the most frequently used of the criteria listed here. Fig1B1 shows a close up of the blebbing region after the event. The point of furthest extent is denoted *c* in this figure.

Another important feature was the shape of the bleb. The bleb in Fig1B1 is mostly a circular arc. When it is circular then the nucleation point should be at the midpoint of the segment joining the bleb shoulder or base points. These are the points at the extent of the bleb. When a feature in the cell geometry prevents the cortex/membrane separation at one side, then the bleb is not circular and the nucleation point is not at the midpoint. Another case when the bleb is not circular occurs when the nucleation is closely followed by a second nucleation.

In the case of Fig1B1, the bleb is nearly circular. However, the prior blebs at the actin structure on the right of *b* prevent extension in that direction. Both of these blebs are visible in Fig1A4. Indeed, the second bleb from the prior frame joins the current bleb. Hence, the current bleb appears to extend to the right. Taking this into account, *a* and *b*, Fig1B1, are the shoulder or base points of the nascent bleb. The furthest extent at *c* lines up well with the midpoint of the segment 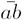.

The third feature as in the old cortex scar in Fig 1B1. We observed a distinct narrow gap. We have denoted this feature *d*. This is likely the means for the cytoplasm to fill the nascent bleb with actin and other proteins sufficient to form a new cortex. This feature, *d*, is about at the midpoint between *a* and *b* and opposite *c*. We frequently used the presence of a gap in the early cortex scar to identify nucleation. As the old actin cortex is disassembled, this sort of indicator will be obliterated.

Another indicator of nucleation is the streaks inside the nascent bleb caused by the moving actin fragments. This feature is an artifact of the slow shutter speed. It is common during the initial frames of a bleb. The streaks indicate the direction of the fluid flow carrying the actin toward the new cell boundary. The source of the flow, the nucleation point must be at the origin of the streaks. In some cases the streaks are the only means to locate nucleation.

When there are multiple nucleation sites, then the streaks display the presence of multiple flows with a turbulent patch between. In this case the furthest extent will indicate which is primary.

It may be impossible to identify a nucleation site. The most common problem was we did not have an image of the bleb at the right time.

In final analysis, for the example all indicators point to *d* as the nucleation site. Note that this conclusion is based solely on observation. Moreover, the example cell is unusual as it is rare to have all these features present for a given bleb.

When the nucleation point was identified, then the next step was to identify the corresponding point on the prior image. Eventually, we applied the energy functional to this location. Even though the cell in general was usually little changed from frame to frame, this may not be true of the region near the bleb.

The first step in this process was to locate points corresponding to the bleb shoulder points. For the current example, we used the cortex features to identify points in the prior state that corresponds to *a* and *b*. We have identified these as *a′* and *b′* in Fig1C. Now, it is clear that the predecessor to *d* is the area we have marked as *d*′.

There is already a visible gap at *d*′. This is not a rare event. In fact we have observed it in over 70% of the blebs we have looked at. Refer to Table 1 line 5.

**Table 1.** Local Combined Prediction Results.

|  | Count | Percentage |
| --- | --- | --- |
| **Total Number of Cells** | 33 |  |
| **Total Number of Blebs** | 142 |  |
| **Confirmed Nucleation from ROI** | 109 | 76.8 % |
| **Predicted by Energy Functional** | 105 | 96.8 % |
| **Confirmed gaps** | 79 | 72.5 % |

An alternate approach was to look at the internal actin structures and GFP bundles. Two such structures are identified in the Fig1B2 for the post bleb state. These structures are identified as *α* and *β*. The same structures are present in the pre-bleb state and denoted *α′* and *β′* in Fig1C. Correlating these structures often the only means to locate the predecessor of an identified nucleation site.

Combining this information, we identified *d′* as the predecessor of the nucleation point.

Next, we executed the discrete energy functional for the membrane segment between *a′* and *b′* in Fig1B1. The values are shown in Fig1D. The horizontal axis shows the point id from the membrane point list while the vertical axis is the energy value. We joined the values with line segments to make it easier to determine the sequential order of the values. We have marked the maximal value with and *x*. The arrowhead in Fig1E identifies the corresponding point on the cell boundary. Note that we are pointing at the area of the gap denoted *d′* in Fig1B1. This is the location we arrived at via observation. Hence, the energy functional and the observation return the same result. Table 1 shows our cumulative results. The data show that the energy functional does an excellent job in predicting the nucleation site when applied to the region defined by the expected bleb shoulder points.

We expect that the energy functional would be as successful with the nearly 25% of cases where no nucleation point could be determined. Indeed, the predicted results in those cases seem to be reasonable.

We have proved a correlation between bleb site selection and edge geometry. The correlation is more conclusive than in prior works. In the next subsection we include cAMP in the discussion on bleb nucleation.

### Global Energy and Blebbing

The natural question is whether the energy functional predicts bleb nucleation when applied to the entire cell. We began by looking at the 10 points with highest energy values for the cell and frame we have been studying. See Fig2A. The points are marked with small dots. The first observation is that the nucleation site we have been studying is not included. This is the one point we know will nucleate at the next frame. Next, many of the high energy locations are at locations where the cell is convex. Bassed on our previous studiy, these are places that are not likely to bleb [27].

Looking closely at the region to the center right, there are 5 high energy locations close together. In Fig2B we plot the energy values for this boundary segment. The energy is high throughout the region. Even though it satisfies geometric requirements for a likely bleb, the underlying biological factors for blebbing are not present. Indeed, this is the current cell posterior. According to [9] Talin concentrations are high in this region. That should suppress the tendency to bleb even though the geometric prerequisite is present.

On the other hand, we restricted attention to the part of the cell boundary where the cAMP gradient influences cell components. In this case, we saw that the next nucleation site ranked high in the energy top ten more than 70% of the time. This is illustrated in Fig2C. This figure shows cumulative results for high energy in the cAMP affected region and bleb nucleation. We see now that the top 3 energy values account for 40% of new blebs and the top 6 for 60%. Furthermore, a Chi-Square test showed that the distribution is significant with 95% confidence. It is stated that a cAMP gradient attracts a list of actin side binding proteins including Myosin II [17]. It seems likely that this protein may play a role both in high energy values and bleb nucleation.

In the following subsection we see that high energy values also indicate high membrane tension. This too will point toward Myosin II as the initiator of blebs as well as the reason for high energy values.

### Comparison of the Energy Functional against Several Alternatives

We considered several alternatives to the energy function. Each time we asked the same question, does the alternative perform as well when asked to predict the next nucleation site given the bleb shoulder points. This is the same question asked in the section Energy and Bleb Nucleation. We looked at a total of 23 cells and 86 blebs.

First we considered the energy functional before the minimization process. This functional was only successful in 74 cases or 86% of the time. We concluded that the unminimized energy functional defined by fixed length linking proteins is not effective.

Next, we isolated each term in the energy functional. In particular, we removed each term one at a time. The result of this study is displayed in Table 2.

**Table 2.**
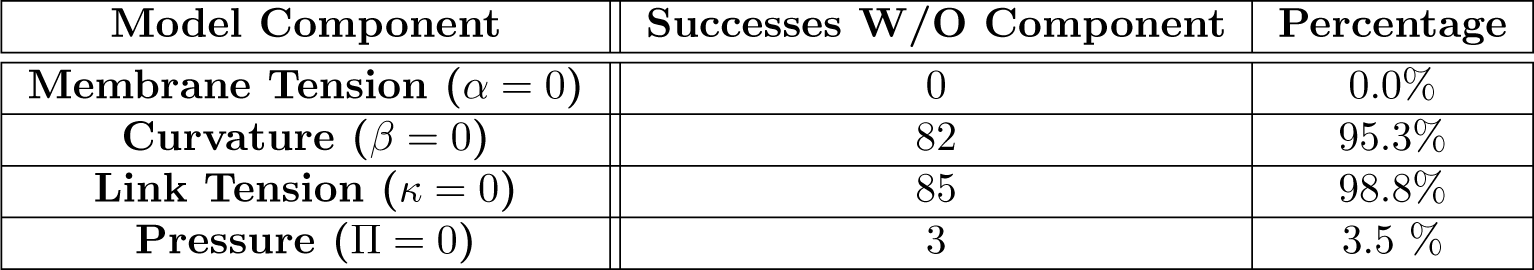
Component Significance within the Energy Function.

| Model Component | Successes W/O Component | Percentage |
| --- | --- | --- |
| Membrane Tension ($\alpha = 0$) | 0 | 0.0% |
| Curvature ($\beta = 0$) | 82 | 95.3% |
| Link Tension ($\kappa = 0$) | 85 | 98.8% |
| Pressure ($\Pi = 0$) | 3 | 3.5 % |

These results are striking. Membrane tension and local pressure are highly significant while curvature and linking protein tension appear to be irrelevant. It would appear that we could remove the latter two terms without changing the effectiveness of the functional. This is something we will need to consider in subsequent realizations of the energy functional. Recall that the pressure term is distinct from the one used in [9] in an attempt to model the local activity of Myosin II. It is likely that both the membrane tension and pressure point toward the presence of Myosin II. This will need to be considered separately as a biological question.

## Conclusion

We have developed an energy functional that predicts the location along the cell membrane where bleb formation initiates. Using data from an under agarose assay, we have shown that our functional accurately predicts over 95 percent of observed bleb nucleation in *Dictyostelium discoideum* as confirmed using cortex gaps, greatest extent, symmetry of the bleb and streak lines in the nascent bleb. This is the case when we restrict attention to the boundary segment between the bleb shoulder points.

The presence of a gap or developing gap at the predicted nucleation site leads us to lean toward cortex degradation as the predecessor to bleb formation. Indeed, we have tabulated more than 70 % of identified nucleation sites show signs of cortex degradation prior to blebbing.

For cases where we cannot identify the nucleation site via observation, the energy functional returns a prediction. We expect it will be as reliable in those cases.

It is an attractive idea that the highest energy locations would cluster about the Myosin II rich posterior face. However, isolating the cell posterior requires microfluidic channels. On the one hand, we know that high energy correlates with membrane tension and we know that the posterior end is rich in Myosin II contracting the cortex. Meanwhile, in [9], Collier et al show that the high Myosin II concentration is offset by high concentration of Talin, which should suppress blebbing. A study that clearly isolate the anterior region will improve global prediction.

There are three elements of this study that separate us from [9], the pressure term in our energy functional is not constant, the underlying geometric platform is based on a B-spline defined from the points returned by the edge detect and we have asked different questions. The result of these changes is a highly successful predictive functional that we believe will point to the origin of blebbing. Furthermore, II = 0 is equivalent to a constant pressure term. The data in Table 2 show clearly that this is not a viable option. Hence, this is a critical extension of its predecessor.

We expect that while the energy framework is highly predictive, it still does not account for all the complex processes influencing the selection of nucleation sites. Aided by our mathematical model, biological experiments can now be designed to study the local mechanism responsible for bleb initiation.

In turn, more biology should be implemented into the energy functional. For instance, Collier et al [9] report the uneven distribution of Talin at the posterior face. To include this information into the energy functional requires developing the functional without equally spaced points, hence without the finite difference method. This is beyond the scope of the current article yet within known numerical processes. This is a matter looking at the several alternatives and determining what adapts best to the case at hand.

## Acknowledgments

The authors thank Chandra Mangroo and Michael Barile for their assistance in data collection. We thank L. Leslie Liu for statistical support. All are at Department of Mathematics and Statistics, Hunter College, CUNY, New York, NY USA.

This work was supported by grants to Derrick Brazill from the National Science Foundation (MCB-1244162), a PSC-CUNY grant (69271 00 47), as well as Research Centers in Minority Institutions Program grants from the National Institute on Minority Health and Health Disparities (8 G12 MD007599) from the National Institutes of Health. Zully Santiago is supported by the Research Inititaive for Scientific Enhancement (RISE) at Hunter College funded by NIH grant GM060665.

## References

1. Blanchoin, L, Boujemaa-Paterski R, Sykes C, Plastino J. Actin dynamics, architecture and mechanics in cell motility. Physical Review, 2014;94:235–263. doi:10.1152/physrev.00018.2013.

2. Blaser H, Reichman-Fried M, Castanon I, Dumstrei K, Marlow F, Kawakami K, Solnica-Krezel L, Heisenberg C, Raz E. Migration of zebrafish primordial germ cells: a role for myosin contraction and cytoplasmic flow. Dev. Cell., 2006;11: 613–627.

3. Cartrina I, Marras S, Bratu D. Tiny Molcular Beacons LNA/2’-O-methyl RNA Chimeric Probes for Imaging Dynamic mRNA Processes in Living Cells. ACS Chem. Biol., 2012;21(9):1 586–1595.

4. Charras G, Yarrow J, Horton M, Mahadevan L, Mitchison T. Non-equiligration of hydrostatic pressure in blebbing cells. Nature Letters 2005;435(19): 365–369.

5. Charras G, Hu C, Coughlin M, Mitchison T. Reassembly of contractile actin cortex in cell blebs. J. Cell Biol., 2006;175: 477–460.

6. Charras G, Paluch E. Blebs lead the way: how to migrate without lamellipodia. Nat Rev Mol Cell Bio., 2008;9: 730–736.

7. Charras G, Coughlin M, Mitchison TJ, Mahedevan L. Life and Times of a Cellular Bleb. Biophysical Journal, 2008 94: 1836–1853.

8. Choi C, Patel H, Barbar D. Expression of Actin-interacting Protein 1 Suppresses Impaired Chemotaxis of *Dictyostelium* Cells Lacking the *Na*+*H*+ Exchanger NHE1. Mol. Bio. Cell, 2010;21: 162–170.

9. Collier S, Paschke P, Kay R, Bretschneider T. Image based modeling of bleb site selection. Scientific Reports, 2017;7: doi:10.1038/s41598-017-06875-9.

10. Crevanna A, Arciniega M, Dupont A, Mizuno N, Kowalska K, Lange O, Wedlich-Soldner R, Lamb D. Side-binding proteins modulate actin filament dynamics. eLife, 2014;e04599 doi:10.7554/eLife.04599.

11. Cunningham C. Actin polymerization and intracellular solvent flow in cell surface blebbing. Journal of Cell Biol., 1995;129: 1589–1599.

12. Evans L. Partial Differential Equations. Amer. Math. Soc,1998.

13. Fackler O, Grosse R. Cell motility through plasma membrane blebbing. J. Cell Biol., 2008;181: 879–884.

14. Dejling H, Helfrisch W. Red blood cell shapes as explained on the basis of curvature elasticity. Biophysical J., 1976;16: 861–868.

15. Friedl P, Wolf K. Tumor-cell invasion and migration: Diversity and escape mechanisms. Nat. Rev. Cancer, 2003;3: 362–374.

16. Helfrich W. newblock Elastic properties of lipid bilayers: theory and possible experiments. newflock Z. Naturforsch., 1973;28: 693–703.

17. Ibo M, Srivastiva V, Robinson D, Gagnon Z. Cell blebbing confined microfluidic environments. PLoS One, 2016; doi:10.1371/journal.pone.0161366.

18. Kaass, M, Witkin A, Terzopoulos D. Snakes: Active Contour Models. Internat. J. Comp. Vision, 1987;119: 3833–3844.

19. Keller H, Eggli P. Protrusive activity, cytoplasmic compartmentalization and restriction rings in locomoting blebbing walker *carcinosarcoma* cells are related to detachment of cortical actin from the plasma membrane. Cell motility and the cytoskeleton, 1998;41: 181–193.

20. Loustau J. Elements of Numerical Analysis with *Mathematica*. World Scientific Press, 2017.

21. Manakiva K, Yan Y, Lowengrub J, Allard J. Cell Surface Mechanochemistry and the Determinants of Bleb Formation, Healing and Travel Velocity. Biophysical J., 2016;110: 1636–1647.

22. Paluch E, Piel M, Prost J, Bornens M, Sykes C. Cortical actomyosin breakage triggers shape oscillations in cells and cell fragments. Biophysics J., 2005;89: 724–733.

23. Pang M, Lynes A, Knecht D. Variables Controlling the Expression Level of Exogenous Genes in *Dictyostelium*. Plasma, 2009;41: 187–197.

24. Pullarkat P. Loss of cell-substrate adhesion leads to periodic shape oscillations in fibroblasts. eprint: arXiv:physics/0612156 2006.

25. Riedl J, Crevenna A, Kessenbrock K, Yu J, Neukirchen D, Bista M, Braedke F, Jenne D, Holak T, Werb Z, Sixth M, Wedlich-Soldner R. Lifeact: A versatile marker to visualize F-Actin. Nat. Methodes, 2008;5(7): 605–607.

26. Sahai E, Marshall C. Differing modes of tumor cell invasion have distinct requirements for Rho/ROCK signaling and extracellular proteolysis. Nat. Cell Biol., 2003;5: 711–719.

27. Santiago Z, Loustau J, Meretzky D, Rawal D, Brazill D. Advances in Geometric Techniques for Analyzing Blebbing in Chemotaxing *Dictyostelium* Cells. PLoS One, 2019, DOI:10.1371/journal.pone.0211975.

28. Strychalski W, Guy R. Computation Model for Bleb Formation. Math. Med. and Biol., 2013;30: 115–130.

29. Strychalski W, Copos C, Lewis O, Guy R. A poroelastic immersed boundary method with applications to cell biology. J. Computational Physics, 2015;282: 77–97.

30. Strychalski W, Guy R. Intracellular Pressure Dynamics in Blebbing Cells. Biophysical J., 2016; 1101168-1179.

31. Sussman M. Cultivation and Syncronous Morphogenesis of *Dictyostekium* under Controlled Experimental Conditions. Methods Cell Biol., 1987;28: 9–29,

32. Tinevez J, Schulze U, Salbreux G, Rosensch J, Joanny, J, Paluch E. Role of cortical tension in bleb growth. PNAS, 2009;106(44): 18581–18586.

33. Trinkaus J. Surface activity and locomotion of Fundulus deep cells during bastula and gastrula stages. Dev Biol., 1979;30: 68–103.

34. Tyson R, Zatulovskiy E, Kay R, Bretschneider T. How blebs and pseudopods cooperate during chemotaxis. PNAS, 2014; 111: 11703–11708.

35. Vellman D, Lemieux M, Knecht D, Insall R. PIP3-dependent macropinocytosis is incompatible with chemotaxis. doi:10.1083/jcb.201309081.

36. Woolley T, Gaffney E, Oliver J, Waters S, Baker R. Global contraction or local growth, bleb shape depends on more than just cell structure. J. of Theoretical Biol., 2015;380: 83–97.

37. Yoshida K, Soldati T. Dissection of amoeboid movement into two mechanically distinct modes. Journal of cell science, 2006;119: 3833–3844, doi:10.1242/jcs.03152.

38. Young J, Mitran S. A numerical model of cellular blebbing: A volume-conserving, fluid-structure interaction model of the entire cell. Journal of Biomechanics, 2010;43: 210–220. doi:10.1016/j.jbiomech.2009.09.025.

39. Zatulovskiy E, Tyson R, Bretschneider T, Kay R. Bleb-driven chemotaxis of Dictyostelium cells. J. Cell Biol., 2014;204(6): 1027–1044, doi:10.1083/jcb.201306147.

